# Monsoonal climate and asymmetric heating facilitate the slope-aspect to influence landscape-scale tree structure in the Western Ghats, India

**DOI:** 10.1101/2024.08.13.607866

**Authors:** Devi Maheshwori, Shreyas Managave, Girish Jathar, Sham Davande

## Abstract

The regional variation of rainfall and length of the rainy season are identified to control the vegetation distribution in the Western Ghats (WG) of India, one of the important biodiversity hotspots. Here we demonstrate the existence of two new modes related to monsoonal climate and asymmetric solar heating that influence the vegetation structure at the landscape-scale in the WG: (i) the north-facing slopes have a higher tree cover (TC) and canopy height (CH) than the south-facing slopes and (ii) higher TC and CH on the west-facing slopes than the east-facing slopes. The asymmetry associated with these modes increases with the slope angle. The interaction of these modes often leads northwest and southeast aspects to have the highest and lowest TC (and CH), respectively. We thus demonstrate that even in low-latitude regions, the slope-aspect plays an important role in determining the TC and CH if the relief is higher. This fundamental driver of vegetation structure needs to be considered while formulating and executing programs aimed at increasing tree cover and conserving biodiversity, especially in the high relief areas in the tropics having seasonal rainfall, such as in the WG.

## 1. Introduction

The Western Ghats (WG) constitutes a sequence of hills running parallel to the west coast of India for about 1600 km (from 8°N to ∼21°N) and covering an area of ∼160000 km^2^ (Das et al., 2006). It is a world heritage site and one of the important biodiversity hotspots having high species endemicity (Myers et al., 2000). Understanding the factors controlling vegetation distribution in the WG is crucial for the conservation of biodiversity and endemic species. The length of the rainy season and total rainfall has been attributed to control the regional scale vegetation distribution in the WG (Champion and Seth 1968; Pascal 1988; Gadgil, 1996; Gimaret and Carpentier, 2003; Nagendra and Ghate, 2003; Prasad et al., 2008; Ramachandra et al., 2016). However, the factors influencing vegetation distribution at landscape-scale within individual protected areas (PAs) are not appreciated.

The slope-aspect is one of the important drivers of vegetation structure at the landscape-scale (Singh, 2018). The effect slope-aspect results in the equator-facing slopes having a relatively warmer, drier microclimate and soil with lower moisture content than the pole-facing slopes (Geroy et al., 2011; Gutiérrez-Jurado et al., 2013; Zhou et al., 2013). This in conjunction with soil edaphic factors leads to the North-facing slopes having a higher vegetation cover than the South-facing slopes in the Northern Hemisphere; the asymmetry being higher on steeper slopes (Smith and Bookhagen, 2021). The north-south asymmetry in the vegetation cover thus established is common to mid-latitude regions (e.g. Champion and Seth 1968; Holland and Steyn, 1975; Badano et al., 2005; Singh, 2018; Smith and Bookhagen, 2021); its influence in the WG, however, is not recognised.

The influence of slope-aspect on the vegetation distribution in the WG was realized only in terms of regional west-east asymmetry in the vegetation distribution. This asymmetry was attributed to the interaction of the west/south-westerly monsoonal winds with the WG which causes higher rainfall in the windward western part (i.e. area lying west of the crest line of the WG hills) than in the leeward eastern part (Fig. 1). This orographic effect controls the regional distribution of the vegetation: evergreen forests on the western side and deciduous forests on the eastern side of the WG (Champion and Seth 1968; Pascal 1988; Gadgil, 1996). The existence of north-south and west-east asymmetries and their interaction in controlling vegetation structure at the landscape-scale is not known. Here we test whether the slope-aspect controls the tree structure, viz. tree cover (TC) and canopy height (CH), at the landscape-scale in the WG.

**Figure 1.**
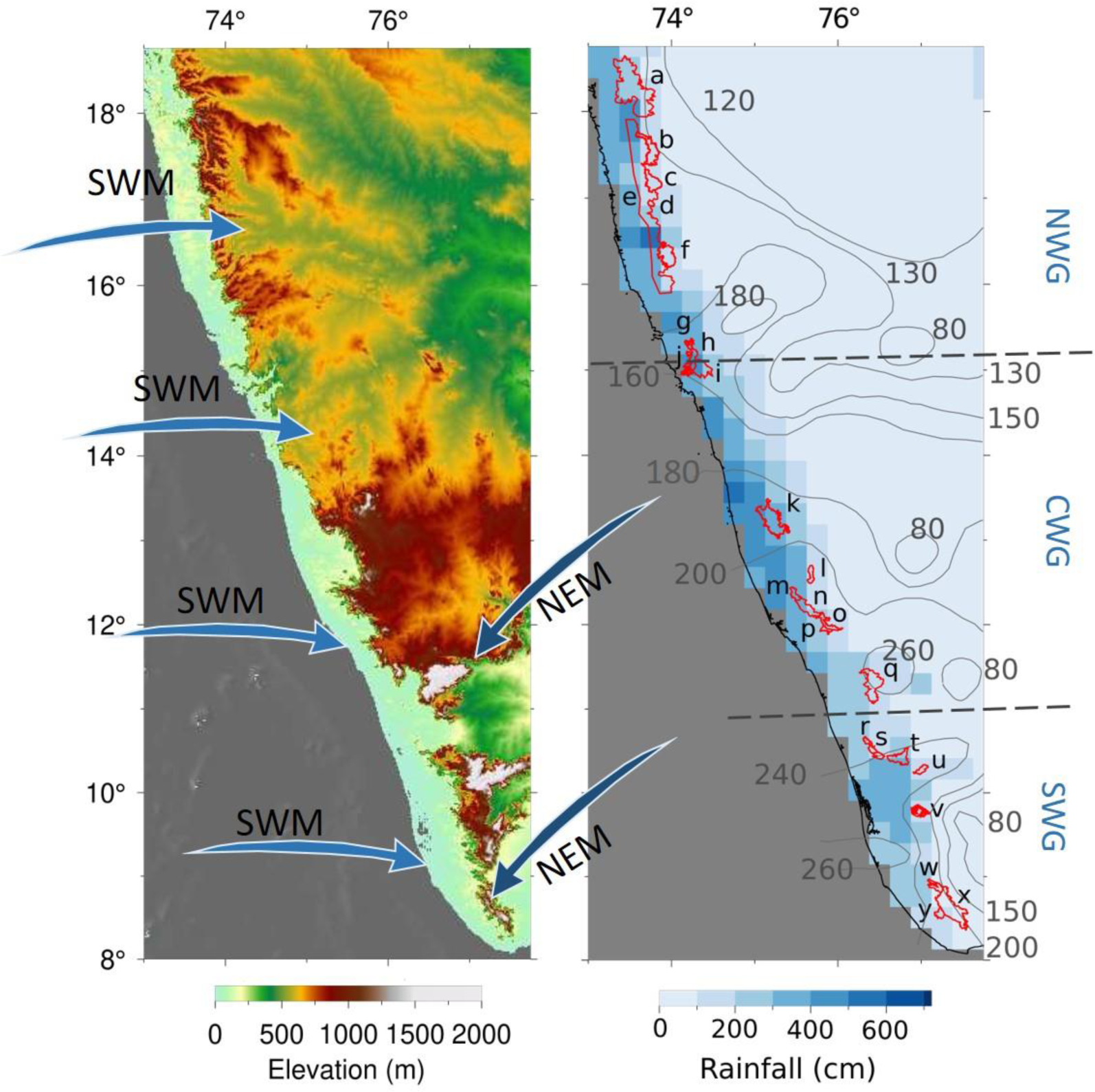
The topography of the WG (based on ASTER Global Digital Elevation Model) (left panel). The arrows indicate the direction of the Low-level winds (925 hPa) (Viswanadhapalli et al., 2020). Distribution of rainfall (Pai et al., 2014) and the length of the rainy season (shown by contours) (Singh et al., 1986) in the WG (right panel). The regions highlighted in red colour (right panel) are protected areas (PAs) analysed for their TC and CH. Northern (NWG), Central (CWG) and Southern (SWG) parts of the WG are demarcated by horizontal dashed lines. The notations for the PAs: a- Tamhini-Lavasa-Raigad; b- Koyna; c-Chandoli; d- Vishalgad; e-‘West of the WG Escarpment’; f-Radhangiri; g-Bhagwan Mahavir; h-Netravali; i-Anshi; j-Cotiago; k-Kudremukh; l-Pushpagiri; m-Talakaveri; n-New Brahmagiri; o- Brahmagiri; p-Aralam; q-New Amarambalam-Silent Valley; r-Peechi; s-Chimmony; t- Parambikulun; u-Indira Gandhi; v-Idukki; w-Shendurney; x-Kallakad Mundanthurai; y- Neyyar.

## 2. Material and Methods

### 2.1. Regional Setting

The WG acts as an orographic barrier to the south-westerly monsoonal winds during southwest monsoon (SWM) season (June to September) which results in higher and lower rainfall on the windward (western margin) and leeward (eastern margin) sides of the WG, respectively (Gunnell, 1997) (Fig. 1). The southern part of WG receives additional rains during the northeast monsoon (NEM) season (October to December) (Fig 1, Suppl. Fig. S1). This, in conjunction with an earlier onset and late withdrawal of SWM, leads the southern part of the WG to have a longer rainy season (∼260 days) as compared to that in the northern part (∼130 days) (Fig. 1) (Singh, 1986).

The seasonal amplitude of temperature is higher in the northern Western Ghats (NWG) than in the central and southern Western Ghats (CWG and SWG, respectively) (Suppl. Fig. S1). In the WG, December and January are the coldest months (∼23⁰C); April and May are the warmest (∼30⁰C in the NWG, and ∼27⁰C in the CWG and SWG) (Suppl. Figure S1). The lapse rate of average annual temperature observed in the southern part of WG varied from 0.5° to 0.7⁰/100 m (von Lengerke,1977). At high altitude PAs (2000-2500 m) the average monthly temperature varies from 9.5 (January) to 18.7 (April) (von Lengerke,1977).

To assess the effect of aspect on TC and CH we studied vegetation in 25 protected areas (PAs) distributed throughout the WG (Fig. 1). This study covers ∼8400 km^2^ of area; different aspect categories cover 9 to 15% of it (Suppl. File 1 and Fig. S2). About ∼62 % of the area in these PAs exhibits a slope between 10° to 30°, followed by 0° to 10° (∼26 %), 30° to 50° (∼11 %) and 50° to 90° (∼1 %). Within individual PAs the area covered by slope 10° to 30° is the most abundant (Suppl. Fig. S3). The median elevation of the PAs varies from 143 to 1346 m; only two PAs (indicated by letters *q* and *u* in Figure 1) have elevations higher than 2000 m. The higher altitudes, especially in the CWG and SWG, were associated with steeper slopes (Suppl. Fig. S4). The high-altitude regions often show a mosaic of trees and grasslands, the shola landscape (Meher-Homji, 1967).

### 2.2. Data sets

To demonstrate the influence of slope-aspect on the tree structure in the WG, we used the Hansen Global Forest Change v1.9 (2000-2021) (Hansen et al., 2013) and Global Forest Canopy Height, GEDI_V27 (Potapov et al., 2021) data to get TC and CH, respectively. Hansen et al., (2013) defined the tree as vegetation taller than 5m in height and tree cover is expressed as a percentage per output grid cell of size 30 x 30 meters. Potapov et al., (2021), who considered the tree as vegetation taller than 3m in height, integrated lidar and Landsat analysis-ready data to produce CH data of 30m resolution. SRTM DEM data (30 m resolution) was used to characterize topography.

### 2.3. Methodology

To avoid the artifacts of anthropogenic activities on the aspect-related asymmetry in vegetation structure, we restricted our analysis to PAs. The boundaries of PAs were taken from The Ministry of Environment, Forest and Climate Change, Government of India’s Decision Support System (MoEF & CC-DSS). The boundaries of PAs Tamhini-Lavasa-Raigad, ‘West of the WG Escarpment’, New Brahmagiri, New Amarambalam-Silent Valley and Vishalgad were selected manually. Further, an area below the altitude of 200m was not considered as it was generally affected by anthropogenic activity. Areas under anthropogenic influences (e.g. plantations and human settlements), deciphered through Google Earth images, were ignored.

We considered the median of the TC and CH on eight aspects i.e. North (N), Northeast (NE), East (E), Southeast (SE), South (S), Southwest (SW), West (W) and Northwest (NW), and three slope angle (0° to 10°, 10° to 30° and 30° to 50°) categories. We did not consider the area having a slope from 50° to 90°, which was less than 1% of the total area. The effect of altitude on TC and CH was assessed for the altitude range of 200-500m, 500-900m, 900-1500m and 1500-2600m. The TC and CH data were overlain on the DEM image in Q-GIS to determine the TC and CH under different aspect, slope and altitude categories.

## 3. Results

The regional distribution of TC and CH were characterised by lower TC and CH in the NWG, and higher TC and CH in CWG and SWG (Suppl. Figure S5). Each PA shows aspect-related variation in the median TC and CH (Figure 2). The range in the median TC and CH on various aspects within individual PAs vary from 2 to 32% and 1 to 8 m, respectively. The distribution of TC (and CH) on different aspects within individual PAs was statistically different (p < 0.05, the two-sample Kolmogorov-Smirnov test).

**Figure 2.**
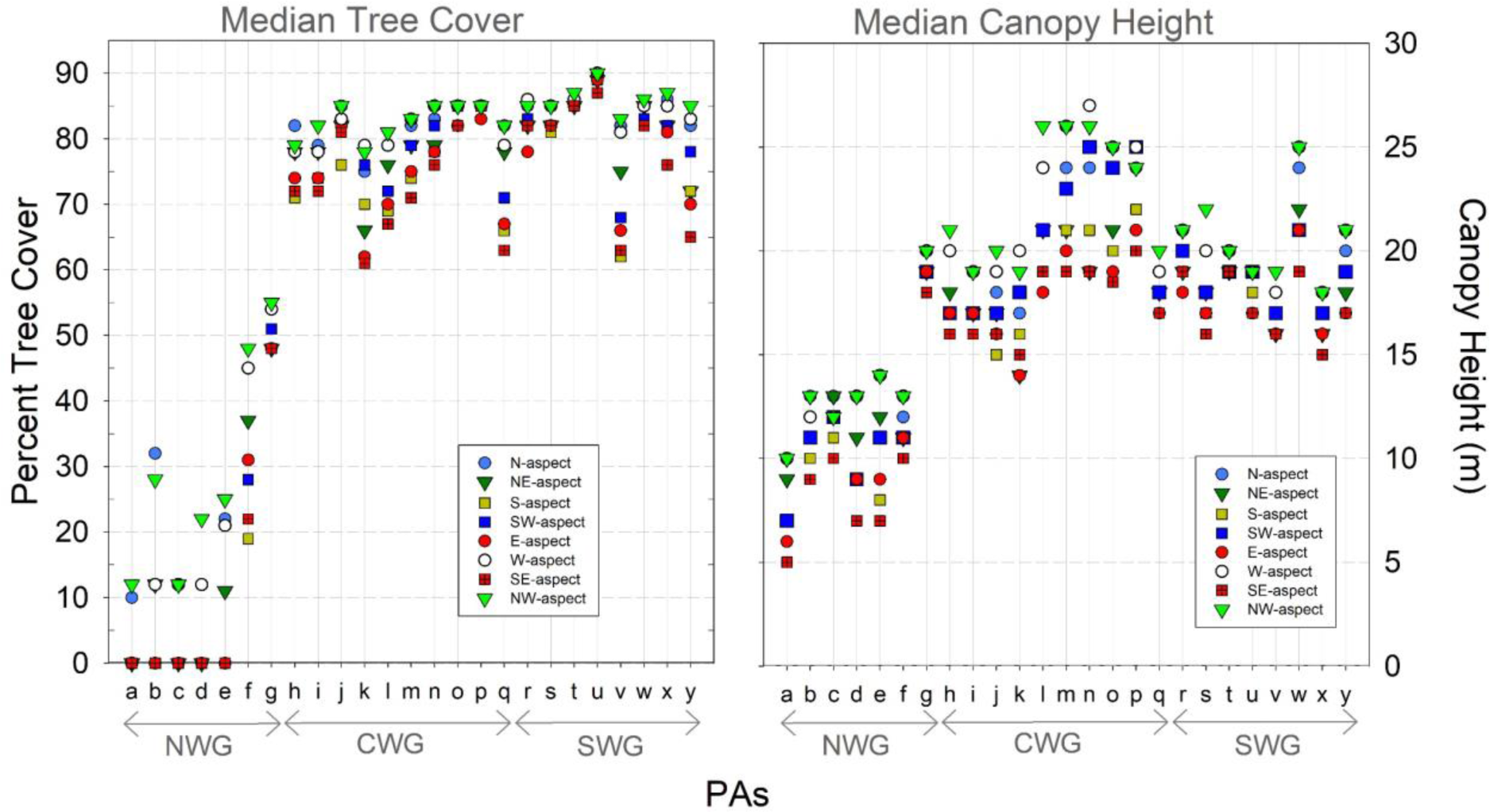
The median tree cover (left panel) and canopy height (right panel) on different aspect categories in the studied PAs. The abbreviations used for the PAs are as in Figure. 1. NWG, CWG and SWG indicate northern, central and southern Western Ghats, respectively.

Even though, the tree was defined differently in TC and CH datasets (i.e. vegetation >5m in TC; >3m in CH) a positive correlation was observed between the median values of TC and CH (i) of different PAs (Suppl. Fig. S5c) and (ii) on different aspects within individual PA (Suppl. Fig. S6).

### 3.1. TC and CH asymmetry on N- and S-aspects

The median TC percent and CH on the N-aspect in the individual PAs is higher than that on the S-aspect (Fig. 3a, b), the N-S asymmetry. This is to say that TC is denser and canopy height taller on the N-aspects than on the S-aspects. Similarly, the median TC and CH showed a decreasing trend from the NE- to E- to SE-aspects (Suppl. Fig. S7) and from NW- to W- to SW-aspects (Suppl. Fig. S8). The N-S asymmetry in the TC (and CH) showed dependence on latitude; although this dependence was statistically significant only for TC (Suppl. Fig. S9).

**Figure 3.**
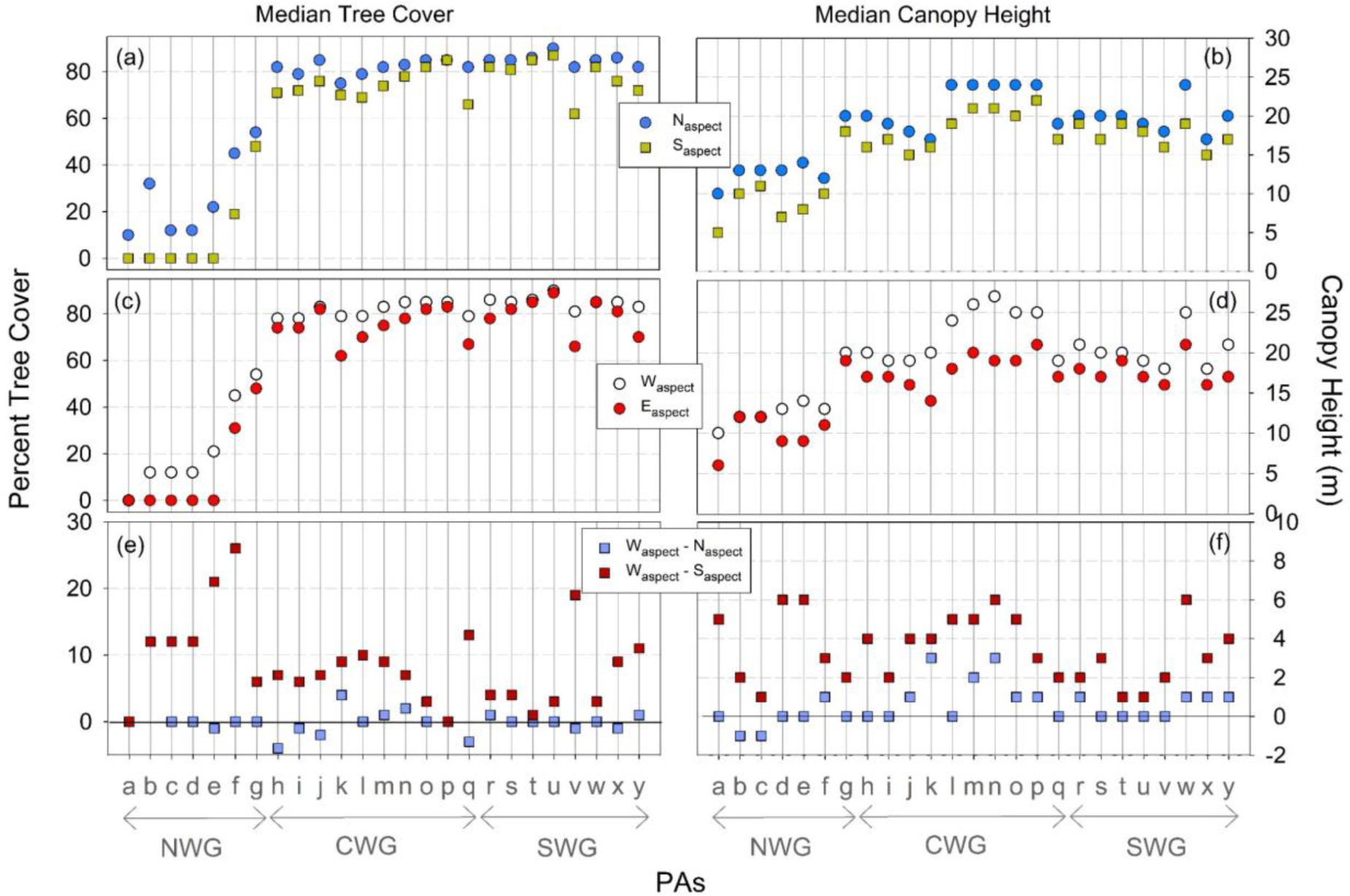
The median tree cover (left panels) and canopy height (right panels) on the N-aspect and S-aspect (a, b) and the W-aspect and E-aspect (c and d) in the studied PAs. The bottom panels show the difference in the tree cover (e) and canopy height (f) on the W-aspect and N-aspect, and on W-aspect and S-aspect. The abbreviations (‘a’ to ‘y’) used for the PAs are as in Figure. 1.

The median TC percent on the N-aspect was similar to that on the W-aspect (Waspect − N_aspect_ = − 1 ± 5%) but the median TC percent on the S-aspect was always lower than on the W-aspect (W_aspect_ − S_aspect_ = 9 ± 6%) (Figure 3e). A similar pattern was also observed for the CH: W_aspect_ − N_aspect_ = 1 ± 1 m and Waspect − S_aspect_ = 3 ± 2 m (Figure 3f).

### 3.2. TC and CH asymmetry on W- and E-aspects

We observed higher median TC percent and CH on W-aspect than on the E-aspect (Fig. 3 c and d) (i.e. W-E asymmetry) which was also manifested in decreasing trend in TC and CH from NW- to N- to NE-aspects (Suppl. Fig. S10) and from SW- to S- to SE-aspects (Suppl. Fig. S11). The correlations of W-E asymmetry in the TC and CH with latitude were statistically insignificant (Suppl. Fig. S9 c and d).

### 3.3. Slope and altitude dependence of the asymmetries in TC and CH

The N-S and W-E asymmetries in the TC and CH increased with the slope angle within individual PAs (Figure 4). The slope-dependent asymmetry is established by occurrence of lower TC and CH selectively on the S- and E-aspects on the steeper slopes (Suppl. Fig. S12 and S13).

**Figure 4.**
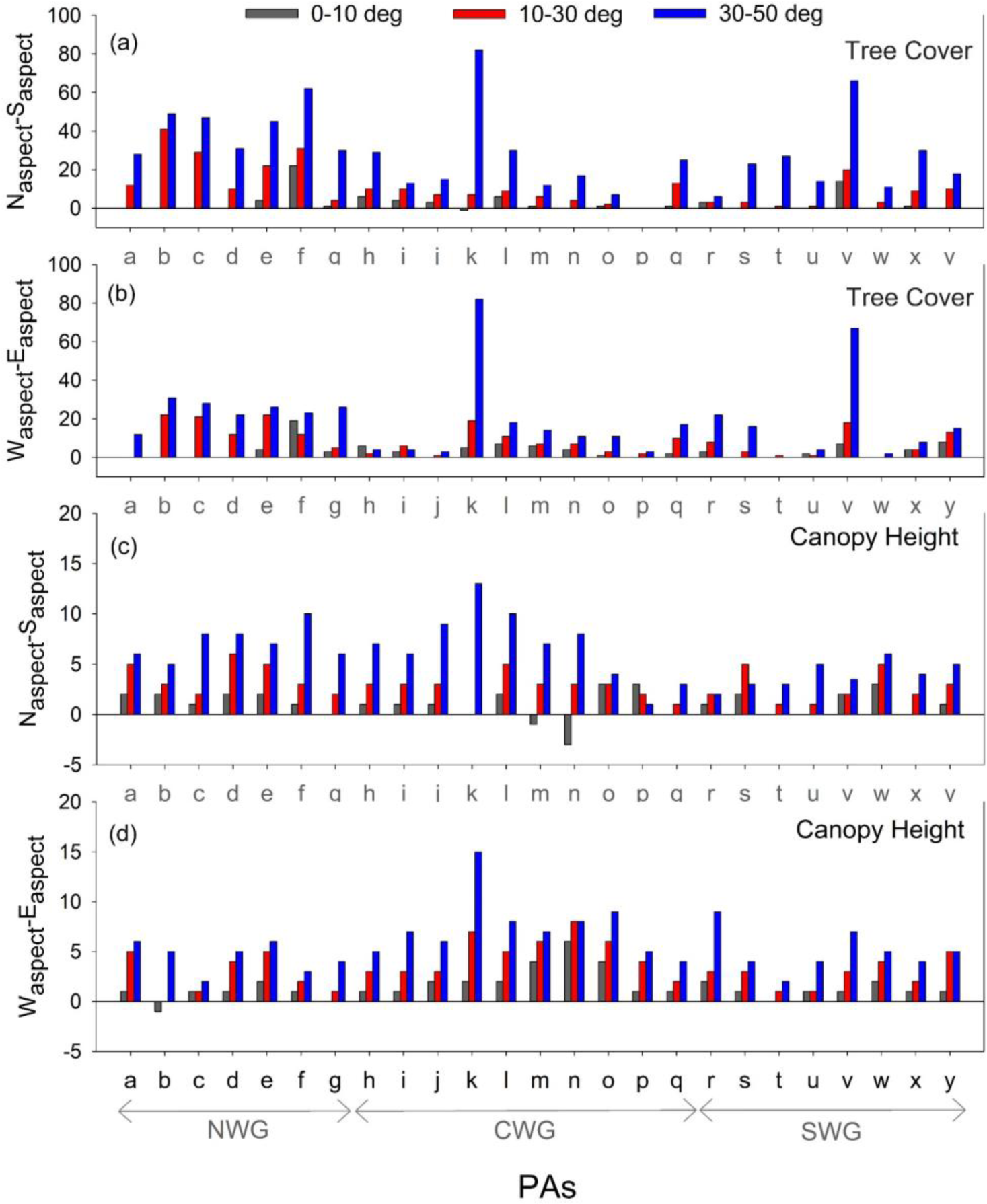
The slope-dependant variation in the difference between the median tree cover on the N-aspect and S-aspect (N_aspect_-S_aspect_) (a) and the W-aspect and E-aspect (W_aspect_-E_aspect_) (b). The same for canopy height is shown in (c) and (d). The abbreviations used for the PAs are as in Figure. 1. Note, the area covered by slope 50-90° is mostly less than 1%.

As compared to their variation with slope, the N-S and W-E asymmetries in TC and CH did not show systematic variations with altitude (Suppl. Fig. S14). Some of the PAs show high values of N-S and W-E asymmetries in TC and CH at 900-1500 m and 1500-2600 m (e.g. PAs *k*, *l*, *n*, *o* and *v* in Suppl. Fig. S14). These altitude bins were associated with steeper slopes. Further, the variations in the TC and CH on N-, S-, W- and E-aspects with altitude (Suppl. Figs. S15 and S16) were more heterogeneous than that with the slope (Suppl. Figs. S12 and S13).

## 4. Discussion

The higher median TC in the CWG and SWG than that in the NWG (Suppl. Fig. S5) reported here is consistent with the earlier finding of the effect of rainfall and the length of the rainy season on the vegetation distribution (Pascal 1988; Ghate et al., 1998; Nagendra and Ghate, 2003; Pascal et al., 2004). Higher rainfall and the length of the rainy season facilitate higher TC in CWG and SWG. The CH is influenced by various factors including hydraulic limitations (Koch et al., 2004; Moles et al., 2009; Tao et al. 2016; Adrah et al., 2022) and sunlight availability (Fricker et al., 2019). The drivers of the CH were found to be scale-dependent: the climate (e.g. water deficit, precipitation) dominates CH at all scales (25 to 1000 m), the relative importance of topography-induced variables (e.g. slope and sunlight exposure) becomes important at fine scale (25-100 m) (Fricker et al., 2019). Here, we observed a positive correlation between TC and CH on coarser (inter-PAs, Suppl. Fig. S5c) and finer (intra-PA, scales between median TC and CH on 8 aspect categories, Suppl. Fig. S6) suggesting similar drivers of TC and CH in the WG. At a landscape scale, the competition for sunlight might have also led the CH to increase where tree cover is higher.

### 4.1. N-S asymmetry: the role of monsoon and solar declination

The TC and CH on the N-aspect were similar to that on the W-aspect (Figure 3 e and f). However, the TC and CH on the S-aspect were consistently less than that on the W-aspect suggesting N-S mode is generated by the factors that selectively lower TC and CH on the S-aspect. This asymmetry was also manifested in decreasing TC and CH from NE- to E- to SE aspects and from NW- to W- to SW-aspects. We attribute the N-S asymmetry to the differential heating of the south-facing and north-facing slopes. We illustrate this with the help of conditions at the northernmost (i.e. Tamhini) and the southernmost (i.e. Neyyar) PAs (Figure 5); the explanation thus also holds for PAs at in-between latitudes.

**Figure 5.**
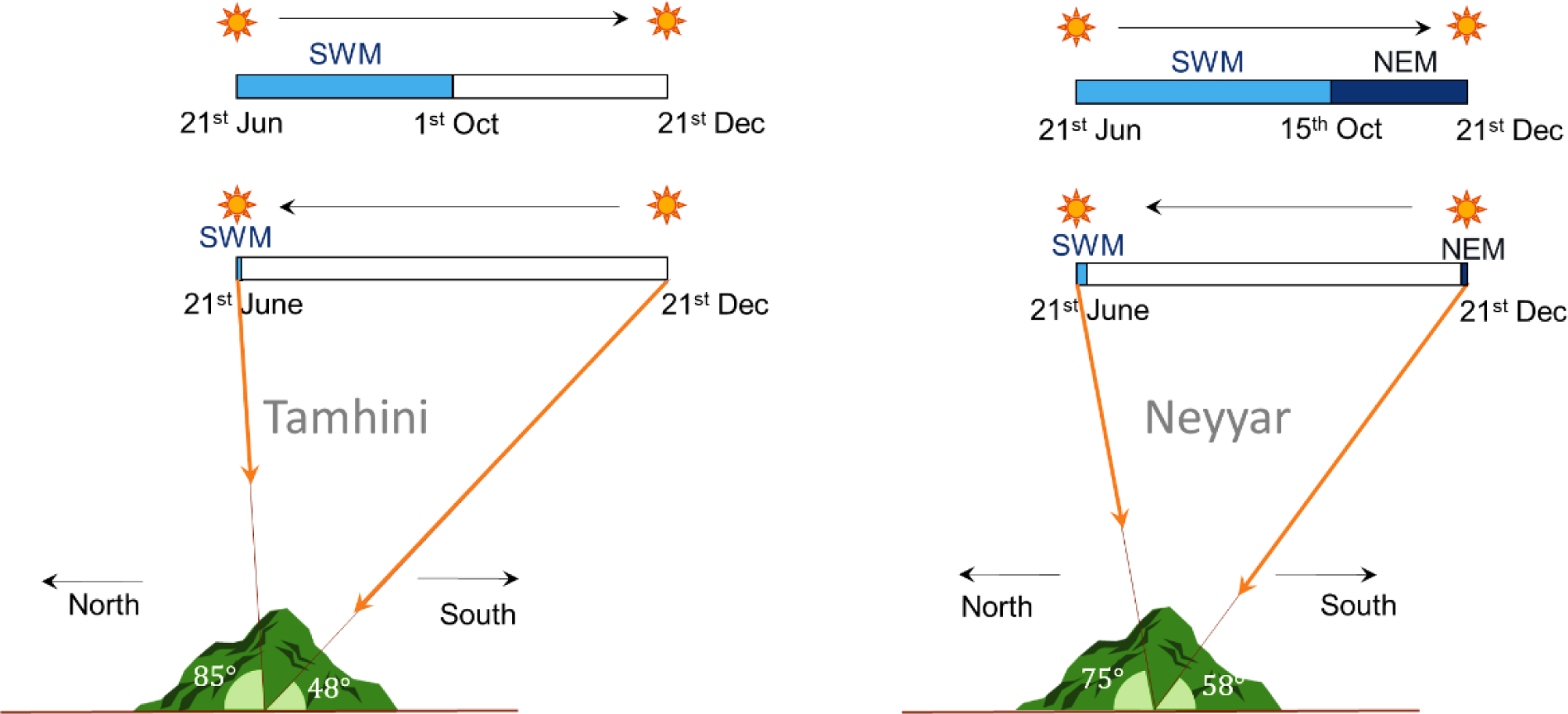
The angle the Sun makes at its highest position (declination) during summer (21st June) and winter (21st December) solstice at Tamhini (the northernmost PA) and Neyyar (the southernmost PA). The figure also shows the SWM and NEM season (shown by shades of blues) vis-à-vis the maximum Sun angles at their onset and withdrawal; the non-rainy season, which is generally cloud-free (India Meteorological Department, 2022) is represented by grey colour. The Sun angle data from www.sunearthtools.com for year 2020.

The duration the Sun spends over the N- and S-aspects is different at these PAs. The Sun is over the S-aspect for 288 days (∼80% of a year, from the 30^th^ of July to the 10^th^ of April) and 225 days (∼62% of a year, from the 1^st^ of September to the 10^th^ of April) at Tamhini and Neyyar, respectively. On the contrary, the Sun is over the N-aspect for a shorter duration at Tamhini (∼20% of a year i.e. for 78 days, from the 12^th^ of May to the 29^th^ of July) and Neyyar (∼38% of a year i.e. for 141days, from the 11^th^ of April to 30^th^ of August).

In addition to the longer duration spend by the Sun over the S-aspect, the monsoonal climate further facilitates higher heating of the S-aspect. Out of the total days the Sun spends over the S-aspect, the non-rainy season (with clearer sky) constitutes 223 days (223 out of 288 days; the 1^st^ of October to the 11^th^ of May) at Tamhini and 100 days (100 out of 225 days; the 1^st^ of January to the 10^th^ of April) at Neyyar. The lesser length of the dry season at Neyyar is due to the northeast monsoon (NEM) rainfall (Figure 5). In comparison, when the Sun is over the N-aspect, only ∼29 (out of 78) and ∼47 (out of 141) days belong to a non-rainy season at these PAs, respectively (Figure 5). Even during these non-rainy days when the Sun is over N-aspect wet and cloudy conditions often prevail due to occurrence the of pre-monsoon showers (India Meteorological Department, 2022). In summary, the south-facing slopes receive sunlight for a significantly longer duration as compared to the north-facing slopes; the asymmetry being higher at Tamhini in the north than at Neyyar located in the south.

The solar radiation load can establish asymmetry in the soil moisture content on north-facing and south-facing slopes. When the Sun is over the southern slopes, the amount of solar radiation received by the southern slopes is higher than the northern slopes (Zhou et al., 2013). In moisture-limited conditions (such as dry season in monsoonal India), the soils on the N-aspect, receiving lower solar heating, have lesser evaporation losses, and thus have more soil moisture content than the soil on the S-aspects (Geroy et al., 2011; Gutiérrez-Jurado et al., 2013; Zhou et al., 2013). The N-aspects also have the potential to retain water for a longer duration (Geroy et al., 2011). The S-aspect, on the contrary, is more soil-moisture deficient (Champion and Seth, 1968; Guan et al., 2015; Lewis et al., 2011).

The vegetation productivity in monsoonal India is mainly controlled by the soil moisture availability (Dubey and Ghosh, 2023), which is the maximum during the rainy season and decreases rapidly afterward; the lowest soil moisture is observed prior to the onset of monsoon (Venkates et al., 2011). Direct sunlight exposure of the southern slopes during the (cloud-free) dry season likely leads to rapid loss of soil moisture on the southern side and can explain the observed N-S asymmetry in the TC and CH in the WG.

### 4.2. W-E asymmetry: an effect of orographic rainfall alone?

The orographic rainfall leads to the evergreen and deciduous forests on the western and eastern slopes of the WG, respectively (Champion and Seth 1968; Pascal 1988; Gadgil, 1996). Thus the conditions are more conducive for tree growth on the western slopes than the eastern slopes. The W-E asymmetry in the TC and CH observed here (Fig. 3, 4) also suggested the influence of the orographic precipitation, especially in PAs that cover the western and eastern slopes of the WG.

The intense orographic rainfall is restricted till 800 m height on the windward side of the WG (Tawade and Singh, 2014); a higher amount of rain was observed on the western slope of the WG and coast plains (Shige et al., 2017; Subrahmanyam and Kumar, 2022). Therefore, the region lying on the western part of the WG crest line is not expected to show W-E asymmetry in the orographic precipitation and hence in TC and CH. However, we observed the W-E asymmetry in TC and CH even over the individual ridges (as low as 400 m in height) lying in the western part of the WG, for example, in ‘West of the WG Escarpment’ and Vishalgad (Figure 1, 2, 6). This alludes to the plausible role of additional factors (such as sea breeze, the persistence of morning dew, and timing of the peak cloud cover vis-à-vis solar inclination) (Smith and Bookhagen, 2021) in establishing W-E asymmetry in the vegetation. In the WG region a higher cloud cover was observed in the afternoon (when the Sun is on the western side) than in the morning (when the Sun is on the eastern side) during non-rainy days (from October to May) (Nikumb et al., 2019). This might facilitate the protection of the soil moisture for a longer duration on the W-aspect post-cessation of the monsoon and lead to the W-aspect having more TC and CH than the E-aspect.

### 4.3. The role of slope, latitude and altitude

The N-S asymmetry in the solar insolation increases with the slope angle and latitude (Smith and Bookhagen, 2021). The slope-dependence of N-S asymmetries in TC and CH (Fig. 4) suggested the role of asymmetric heating of North and South facing aspects in establishing N-S asymmetry in the vegetation structure in the WG. The increase in the N-S asymmetries in TC (and to some extent CH) with latitude (Suppl. Fig. 9a, b) further supports this conclusion.

The orographic rainfall is restricted to the windward side of the WG. A long and gentle rise in topography is associated with a higher orographic rainfall than with a steep rise (Tawade and Singh, 2014). Therefore, an increase in W-E asymmetry in rainfall with slope is unexpected, and less likely to be attributed to the slope-dependent increase in W-E asymmetry in the TC and CH. The slope-dependence of W-E asymmetry in TC and CH is consistent with the effect of cloud-mediated asymmetric heating of W- and E-aspects (explained earlier). The W-E asymmetry is established by lower TC and CH selectively on the E-aspect (Suppl. Fig. S12, 13); steeper E-aspects get heated more in the morning (which are generally cloud-free), leading to higher soil moisture loss and resulting in its lower TC and CH. Persistence of dew for a longer duration (due to lower heating) on the western aspects (Smith, 1978) might also help in retaining higher soil moisture on the W-aspect and support higher TC and CH. As (i) there is no W-E asymmetry in solar insolation (i.e. radiation in the absence of cloud cover) (Smith and Bookhagen, 2021) and (ii) cloud-mediated asymmetric heating of W- and E-aspects is not known to vary with latitude, the W-E asymmetry in TC and CH did not show latitudinal dependence (Suppl. Fig. 9c, d).

The vegetation in WG showed N-S and W-E asymmetry in TC and CH at all altitude bins; the extent of asymmetry, however, did not change systematically with altitude (Suppl. Fig. S14). It thus appears that lapse rate controlled temperature change may not be controlling the magnitude of N-S and W-E asymmetry in TC and CH in the WG. At lower altitudes (<1500 m), the variation in N-S and W-E asymmetry could be due to the effect of precipitation and slope. While the western side receives orographic precipitation, the precipitation decreases with altitude on the eastern side (Figure 1). This makes it difficult to assess whether the variation in the W-E asymmetry with altitude reported here is the effect of altitude or precipitation. In addition, the slope increases with altitude in the CWG and SWG (Suppl. Fig. S4). As the N-S and W-E asymmetry increase systematically with slope (Fig. 4), part of the altitude-induced variation in vegetation asymmetry could also be due to the effect of slope. In the higher altitude regions (>∼1500 m), especially in the CWG, the vegetation is characterized by a mosaic of grassland and shola (i.e. stunted evergreen forest) (Meher-Homji, 1967). The highest N-S and W-E asymmetry in TC and CH in such areas (Suppl. Fig. S14) is due to this “bi-modal” vegetation pattern.

### 4.4. Combined effect of N-S and W-E asymmetries in the TC and CH

Our work indicated two modes control the landscape-scale TC and CH in the WG: the solar declination induced and monsoonal climate facilitated asymmetry in vegetation on N- and S-aspects; and orographic rainfall, and plausibly solar heating, induced W-E asymmetry. These asymmetries can also be observed within the small watersheds. For example, hill ranges within the watershed at Vishalgad in NWG (area ∼41 km^2^) (Fig. 6). The combined effects of the N-S and W-E decreasing trends in vegetation resulted generally in N- and NW-aspects having the highest TC and CH and S- and SE-aspects, the least (Figure 2, 7). Further, as the N-S and W-E asymmetries increase with slope angle, the vegetation on the steeper slopes on S- and SE-aspects exhibit the lowest TC and CH.

**Figure 6.**
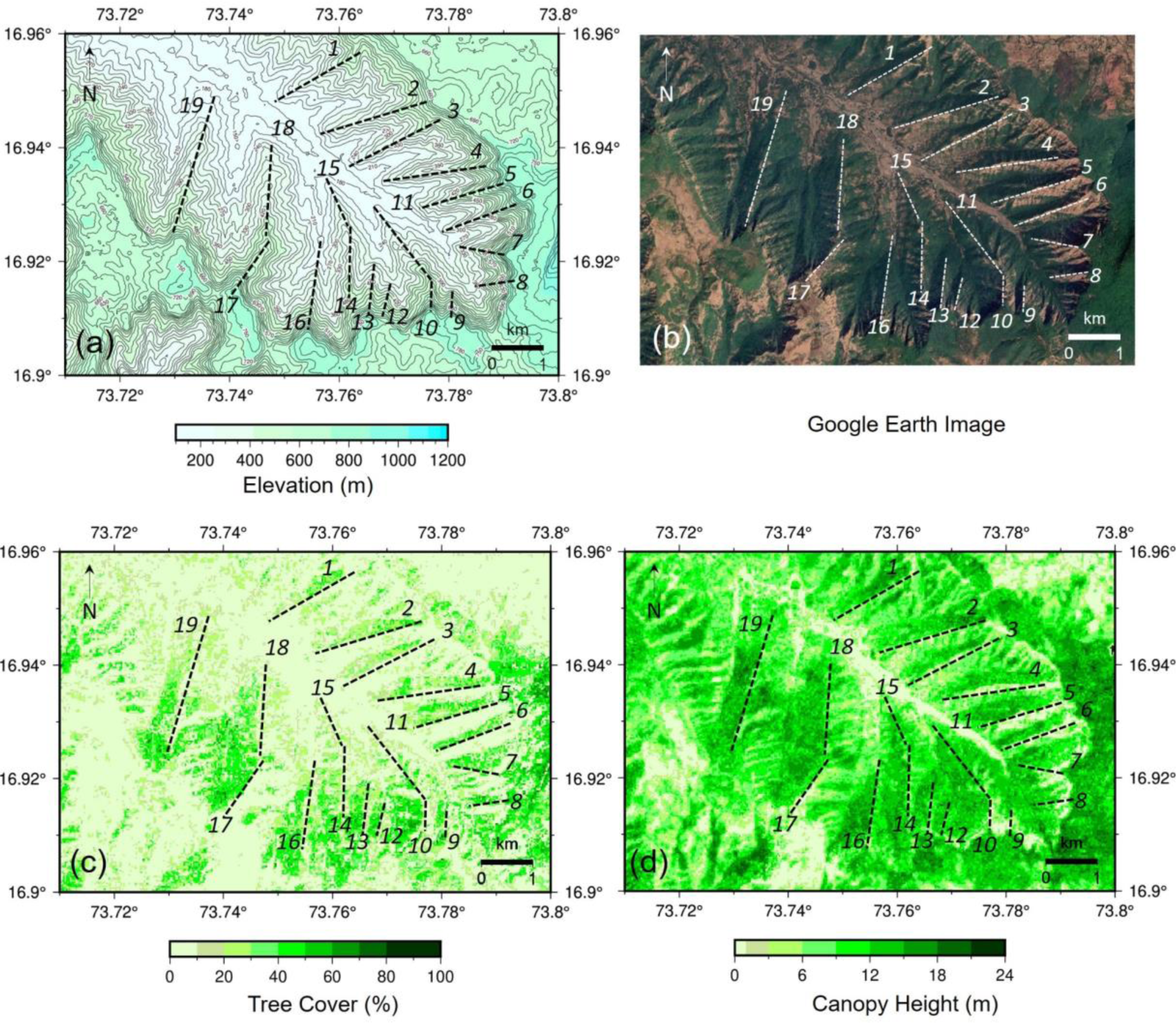
Elevation (a), Google Earth Image (b), Tree cover (c) and Canopy Height (d) at Vishalgad watershed. The dotted lines with numbers indicate ridges.

**Figure 7.**
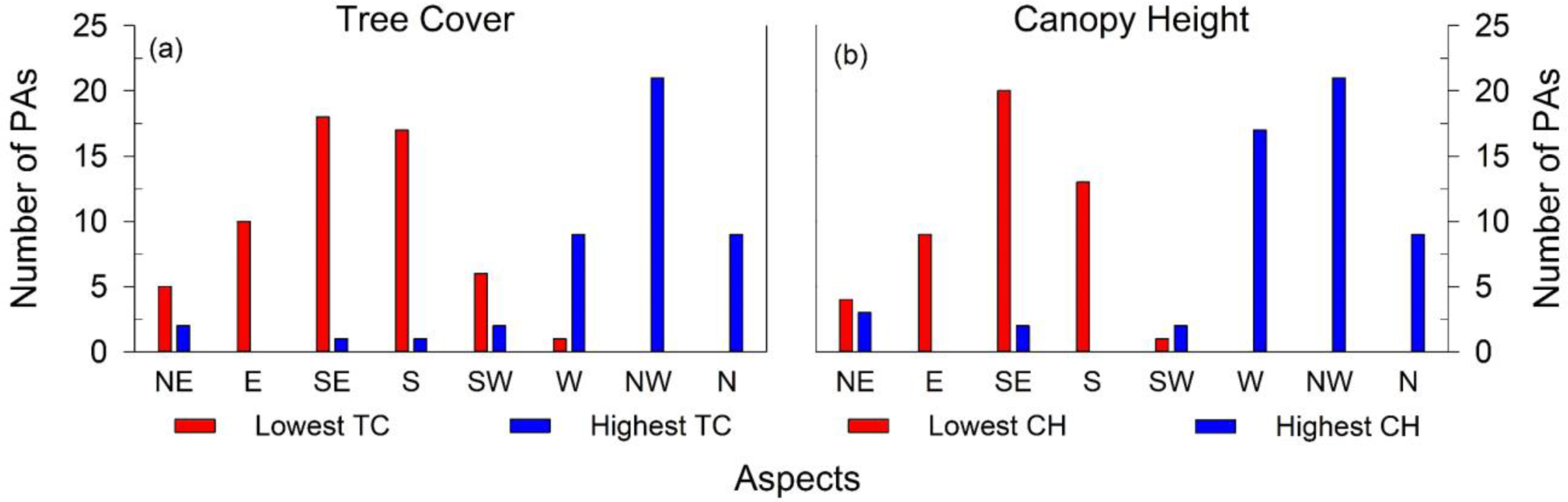
Number of PAs showing the lowest and highest tree cover (a) and canopy height (b) in different aspect categories.

## 5. Conclusion

The WG is an ecologically sensitive zone with high biodiversity and species endemicity (Myers et al., 2000). Understanding the factors controlling vegetation distribution is important not only for academic pursuits but also for the effective implementation of afforestation programs and the conservation of biodiversity. This work demonstrates the slope-aspect influence TC and CH at the landscape-scale in the WG. There is a need to acknowledge the tendency of the southern aspects to have a lower TC and CH as a natural phenomenon; lower TC and CH on the southern aspects should not be considered as “degraded” and a higher TC and CH on northern aspects as “pristine”. The relatively less favourable conditions for tree growth on the S- and SE-aspects should be considered while designing afforestation programs in the WG. Biodiversity assessment studies in the WG should consider the slope-aspect effect as an additional framework while assessing species distribution at the landscape scale. This work shows that slope-aspect can influence the vegetation structure even in the low-latitude regions. This fundamental property of vegetation control needs to be considered as the driver of vegetation structure in the high-relief areas in the tropics, such as in the WG.

## Acknowledgement

We thank Vishwas Kale, Sagar Pandit and Deepak Barua for going through the manuscript and offering suggestion. Help extended by Rahul Tak with programming is appreciated.

## Author contributions

Conceptualization: Shreyas Managave

Methodology: Devi Maheshwori, Shreyas Managave, Girish Jathar, Sham Davande

Formal Analysis: Devi Maheshwori, Shreyas Managave, Girish Jathar, Sham Davande

Investigation: Devi Maheshwori, Shreyas Managave, Girish Jathar, Sham Davande

Resources: Shreyas Managave

Writing—Original Draft: Devi Maheshwori, Shreyas Managave

Writing—Review and Editing: Devi Maheshwori, Shreyas Managave, Girish Jathar, Sham Davande

Supervision: Shreyas Managave

Funding Acquisition: Shreyas Managave

## Materials and correspondence

## Supplementary Information

**Supplementary Figure S1.**
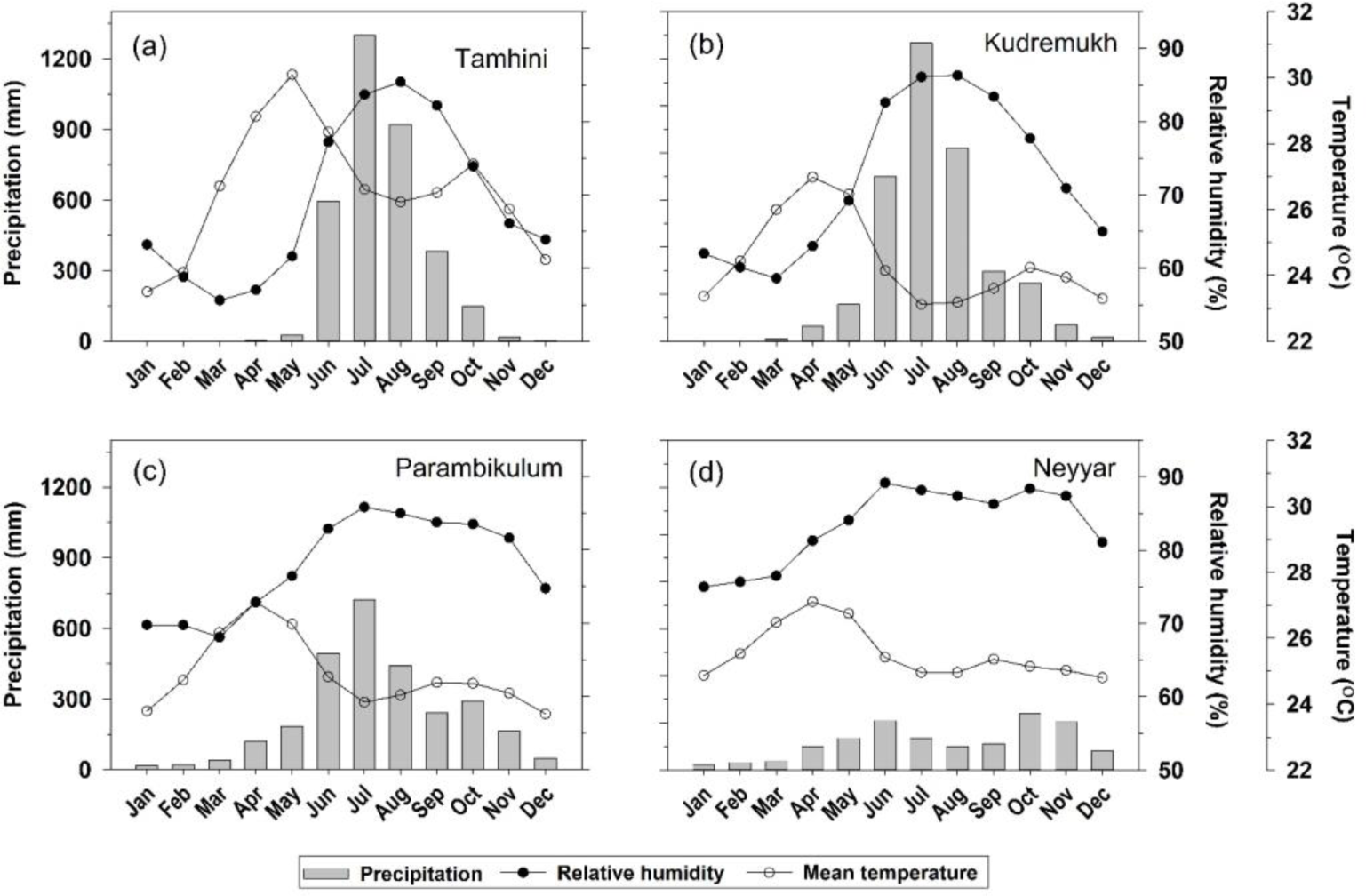
Climatology of monthly precipitation, temperature and relative humidity at selected PAs (a) Tamhini, (b) Kudremukh, (c) Parambikulam, (d) Neyyar. Tamhni is in the NWG, Kudremukh is in the CWG, Parambikulam and Neyyar are in the SWG. NWG, CWG and SWG indicate northern, central and southern Western Ghats, respectively. Data: Harris et al., (2020).

**Supplementary Figure S2.**
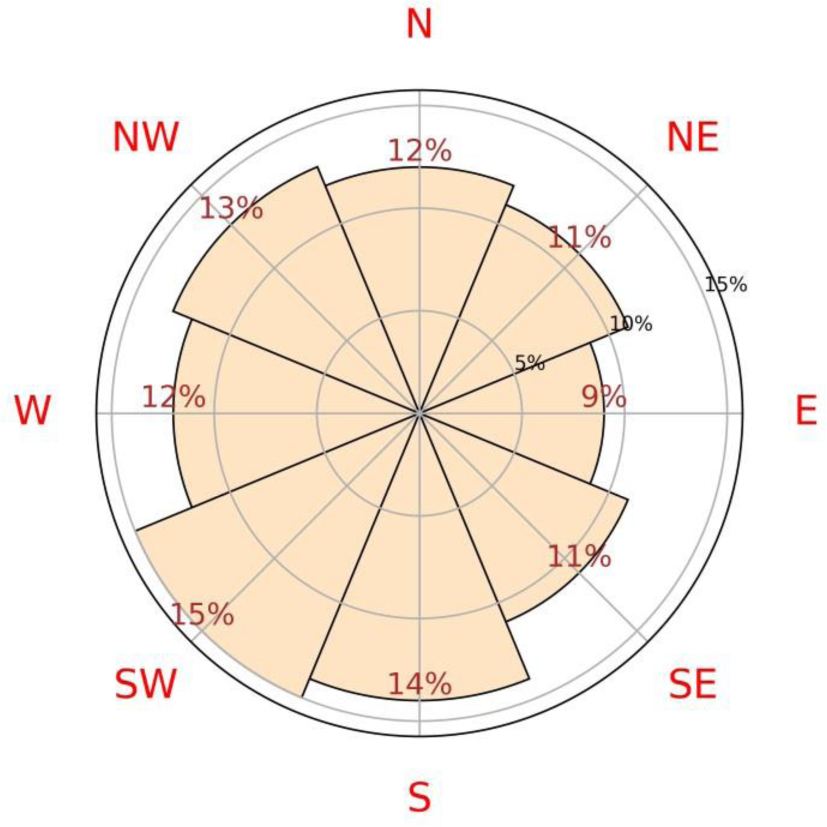
The percent area covered by different aspect categories in the selected PAs. The flat area covers less than 0.05 %.

**Supplementary Figure S3.**
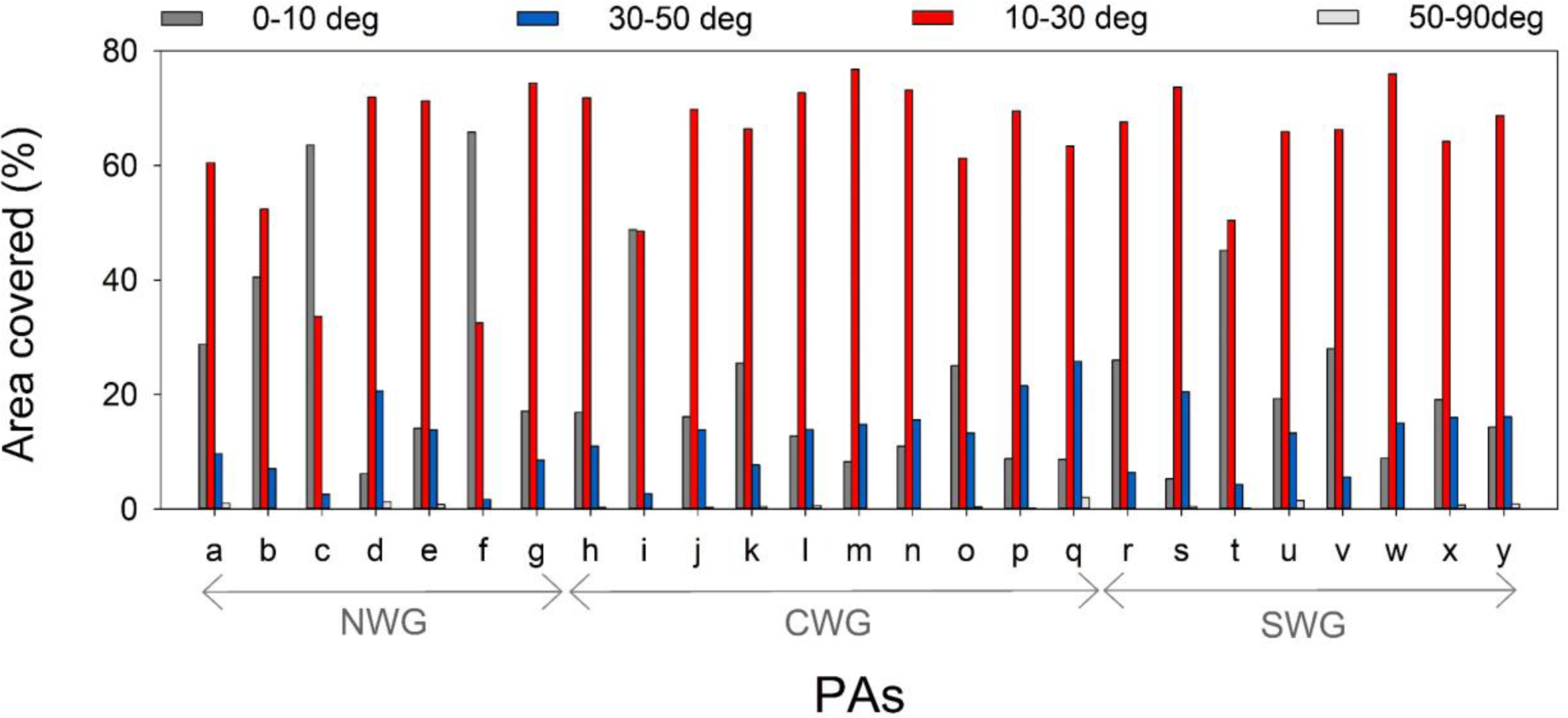
The area covered by different slope categories in the selected PAs. Note, the area covered by slope 50-90° is typically less than 1%. The notations for the PAs: a- Tamhini-Lavasa-Raigad; b- Koyna; c-Chandoli; d-Vishalgad; e-‘West of the WG Escarpment’; f-Radhangiri; g-Bhagwan Mahavir; h-Netravali; i-Anshi; j-Cotiago; k- Kudremukh; l-Pushpagiri; m-Talakaveri; n-New Brahmagiri; o-Brahmagiri; p-Aralam; q- New Amarambalam-Silent Valley; r-Peechi; s-Chimmony; t-Parambikulun; u-Indira Gandhi; v-Idukki; w-Shendurney; x-Kallakad Mundanthurai; y-Neyyar.

**Supplementary Figure S4.**
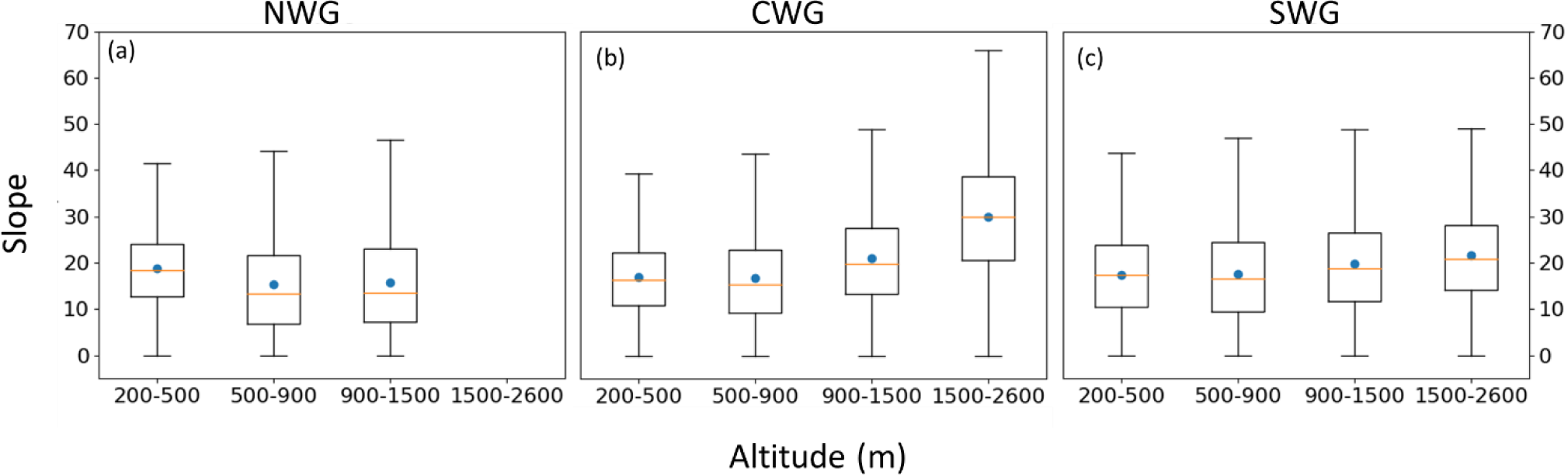
Whisker plot showing variation in the slope in different altitude bands in the NWG (a), CWG (b) and SWG (c).

**Supplementary Figure S5.**
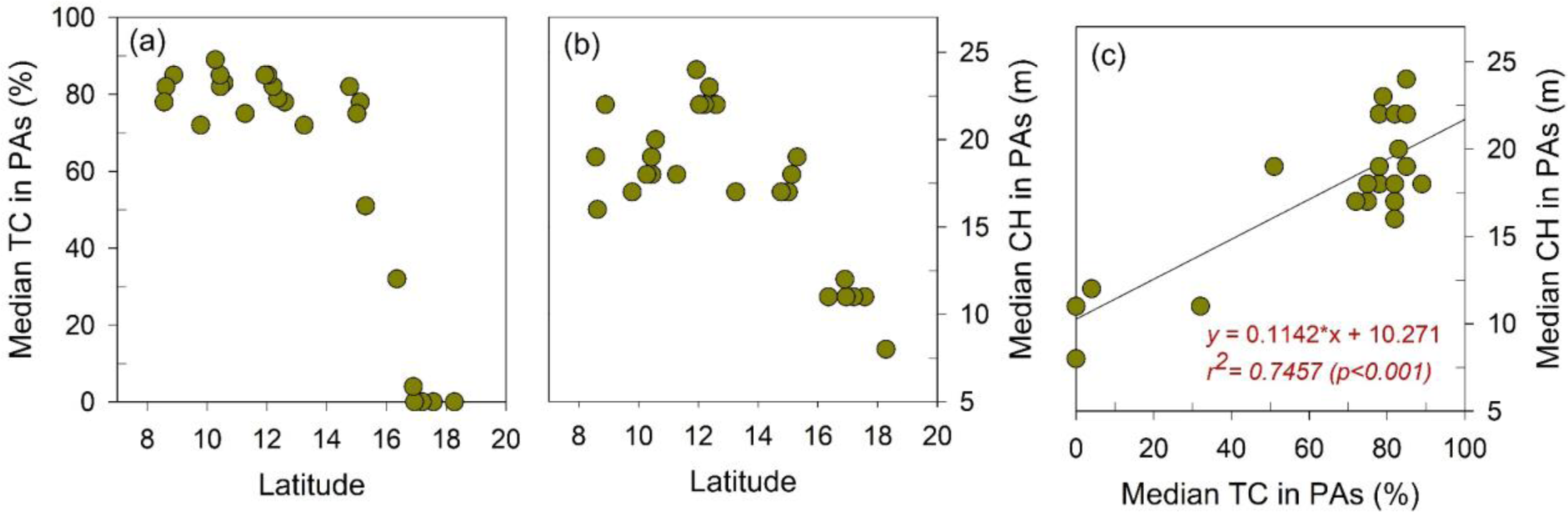
Median tree cover (TC) (a), canopy height (CH) (b) and variation between them (c) in the PAs in the WG.

**Supplementary Figure S6.**
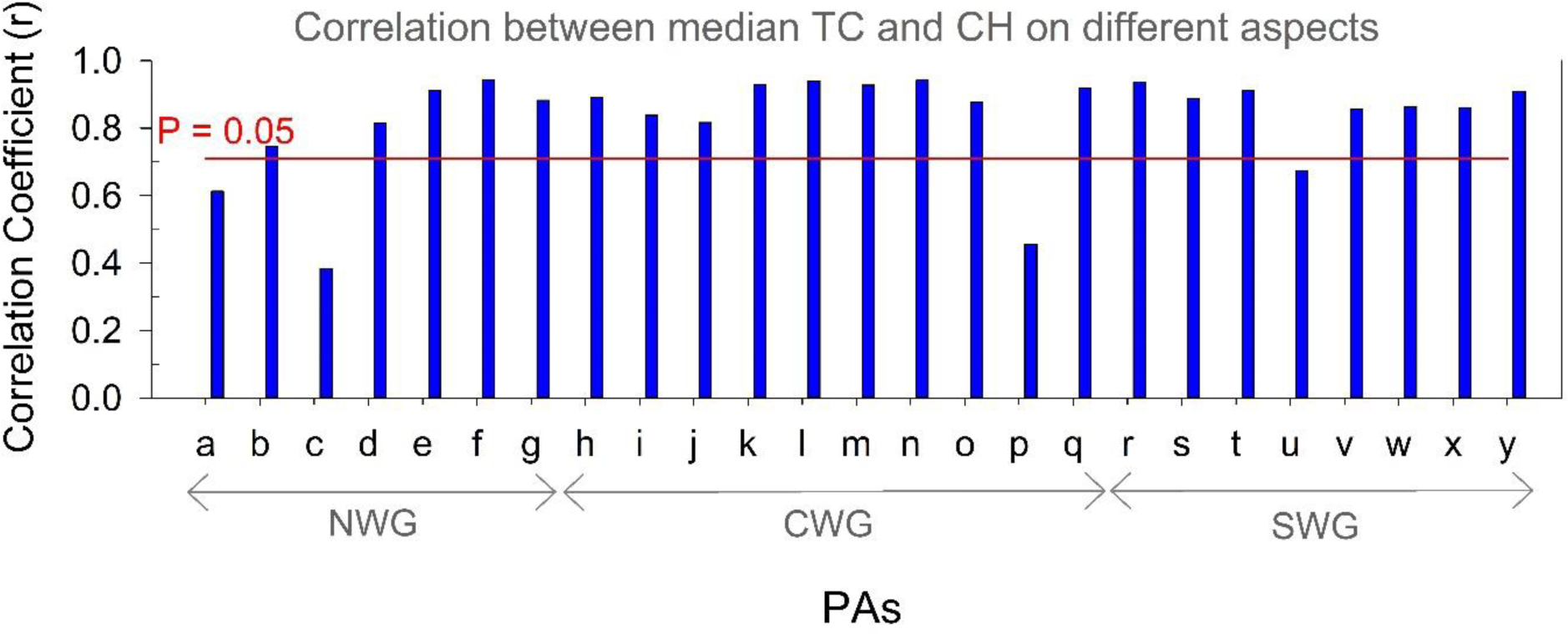
The correlation between the median TC and CH on 8 aspects within PAs. The abbreviations for the PAs are as in Supplementary Figure S3. The horizontal line with ‘P=0.05’ indicates the correlation coefficient at p-value of 0.05.

**Supplementary Figure S7.**
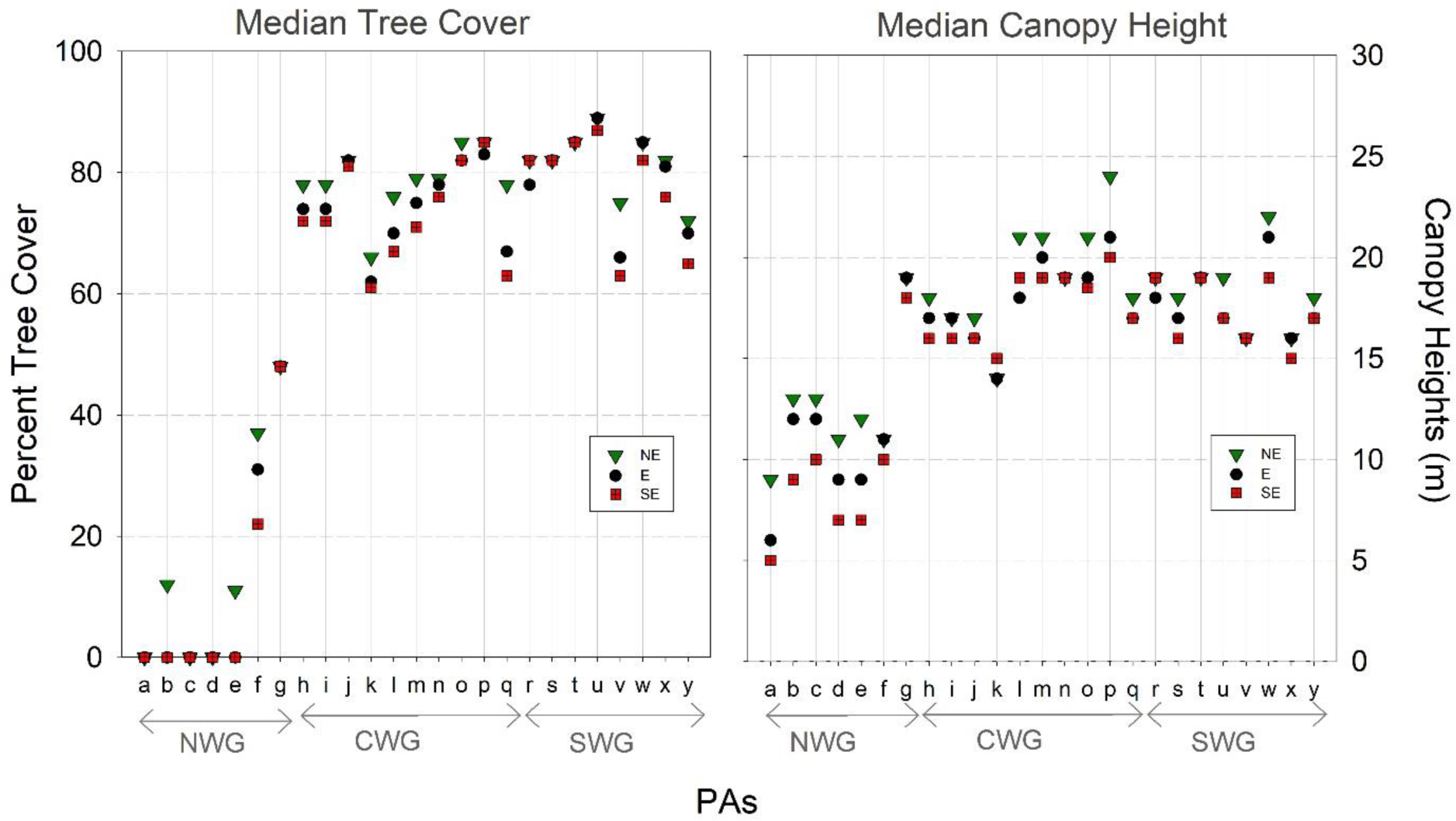
The median TC (left panel) and CH (right panel) on the northeast (NE), east (E) and southeast (SE) aspects in the studied PAs. The abbreviations for the PAs are as in Supplementary Figure S3.

**Supplementary Figure S8.**
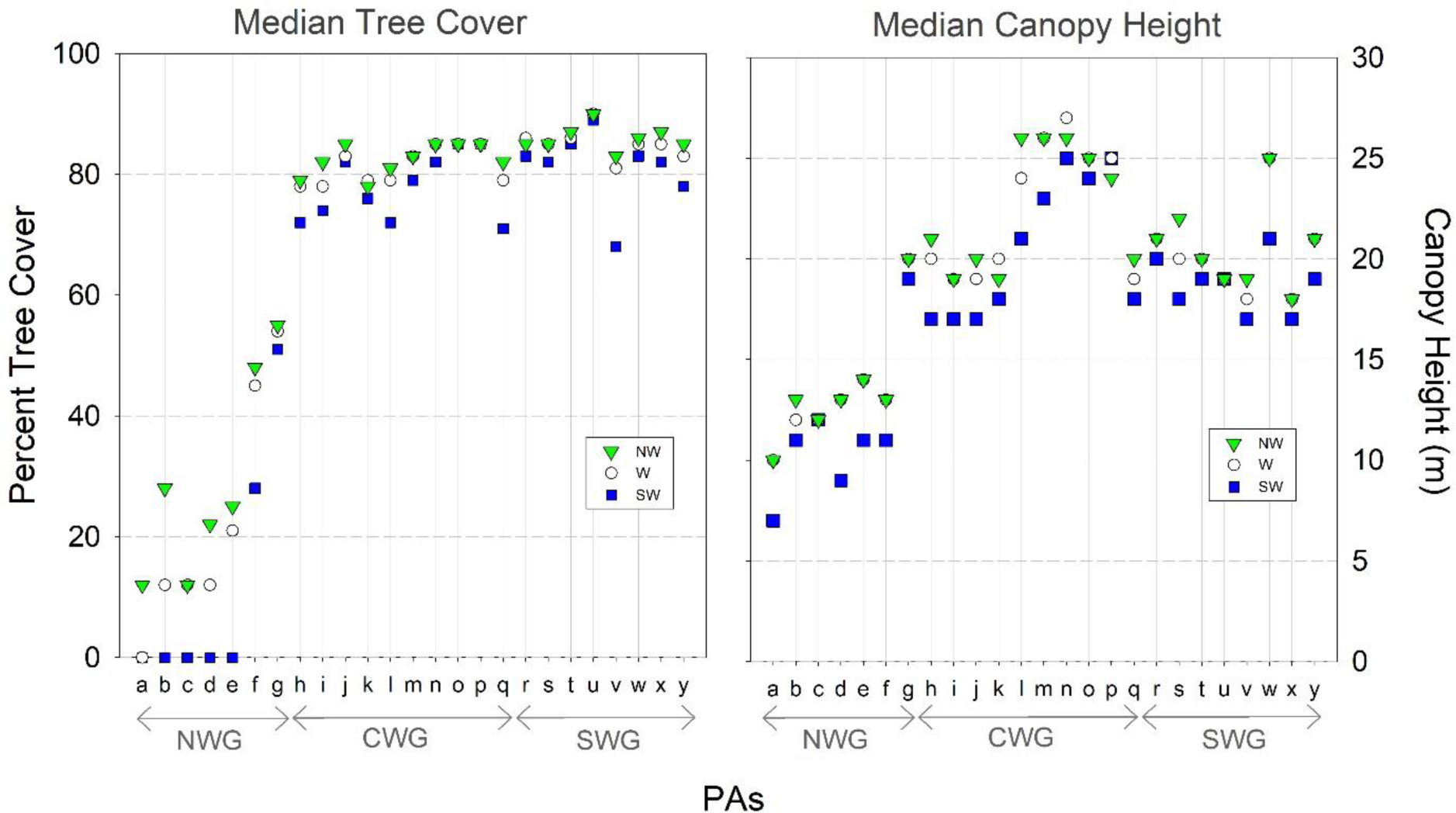
The median TC (left panel) and CH (right panel) on the northwest (NW), west (W) and southwest (SW) aspects in the studied PAs. The abbreviations for the PAs are as in Supplementary Figure S3.

**Supplementary Figure S9.**
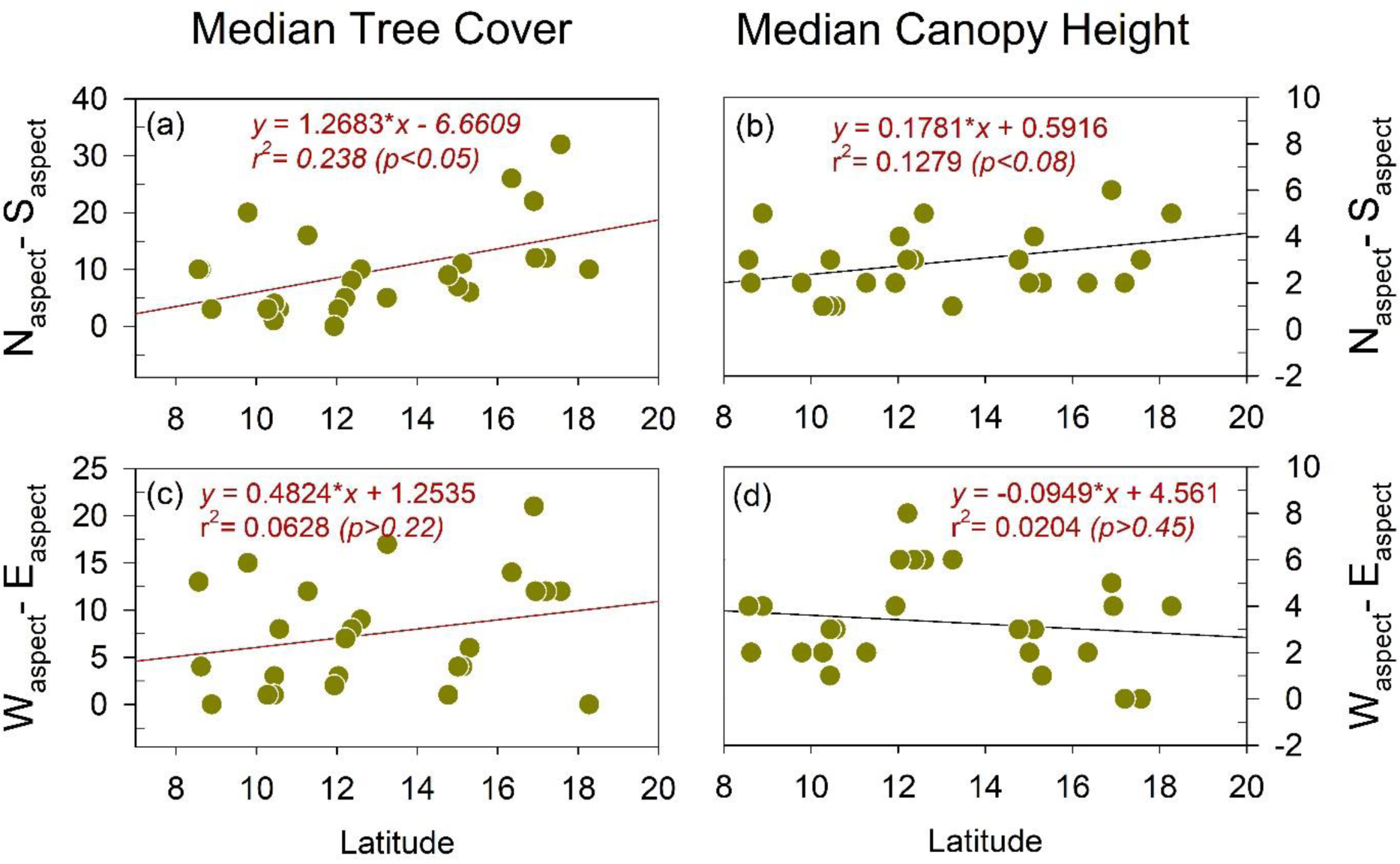
The latitude-dependence of the N-S asymmetry in the TC (a) and CH (b) and of W-E asymmetry in the TC (c) and CH (d).

**Supplementary Figure S10.**
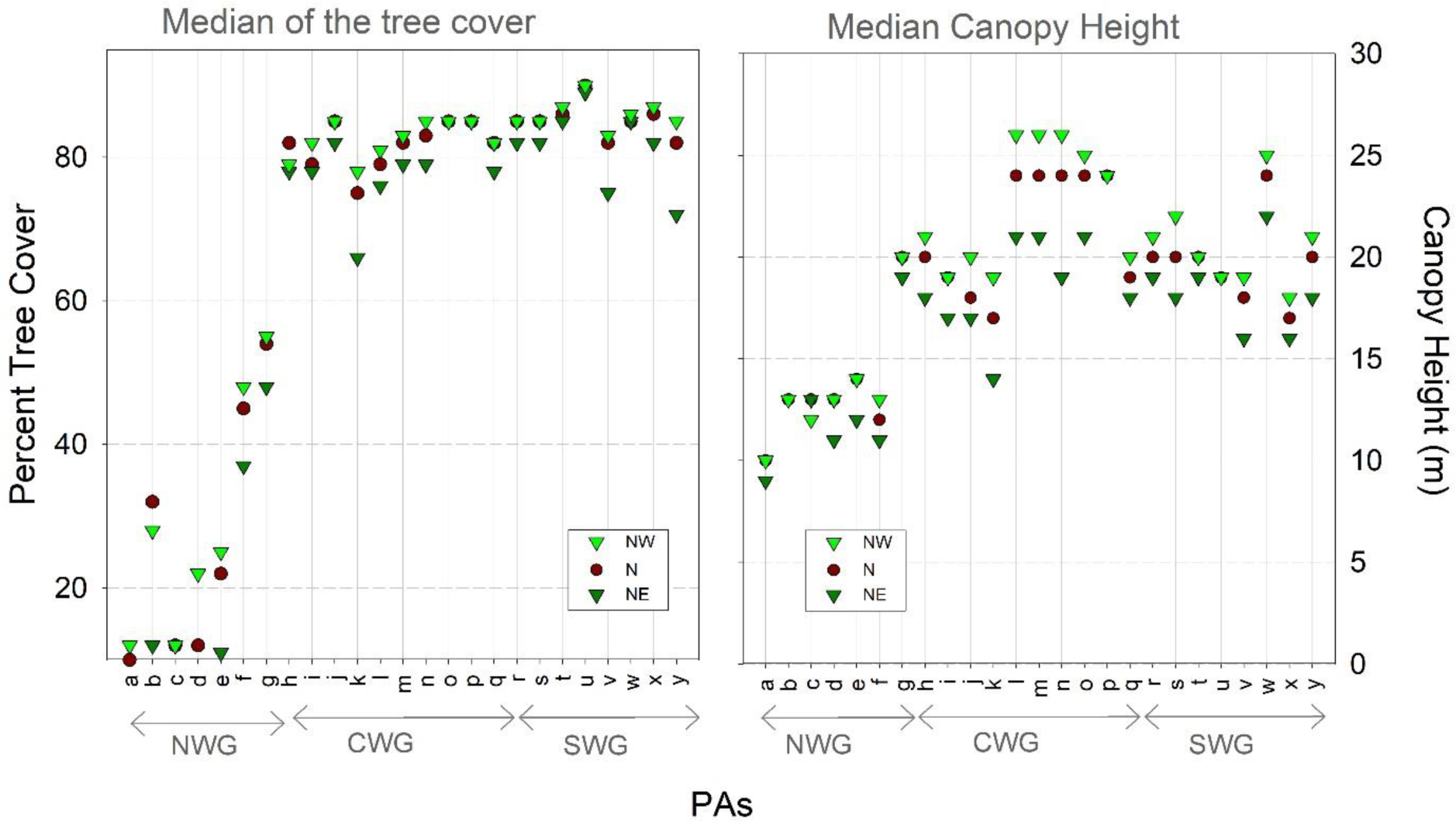
The median TC (left panel) and CH (right panel) on the northwest (NW), north (N) and northeast (NE) aspects in the studied PAs. The abbreviations for the PAs are as in Supplementary Figure S3.

**Supplementary Figure S11.**
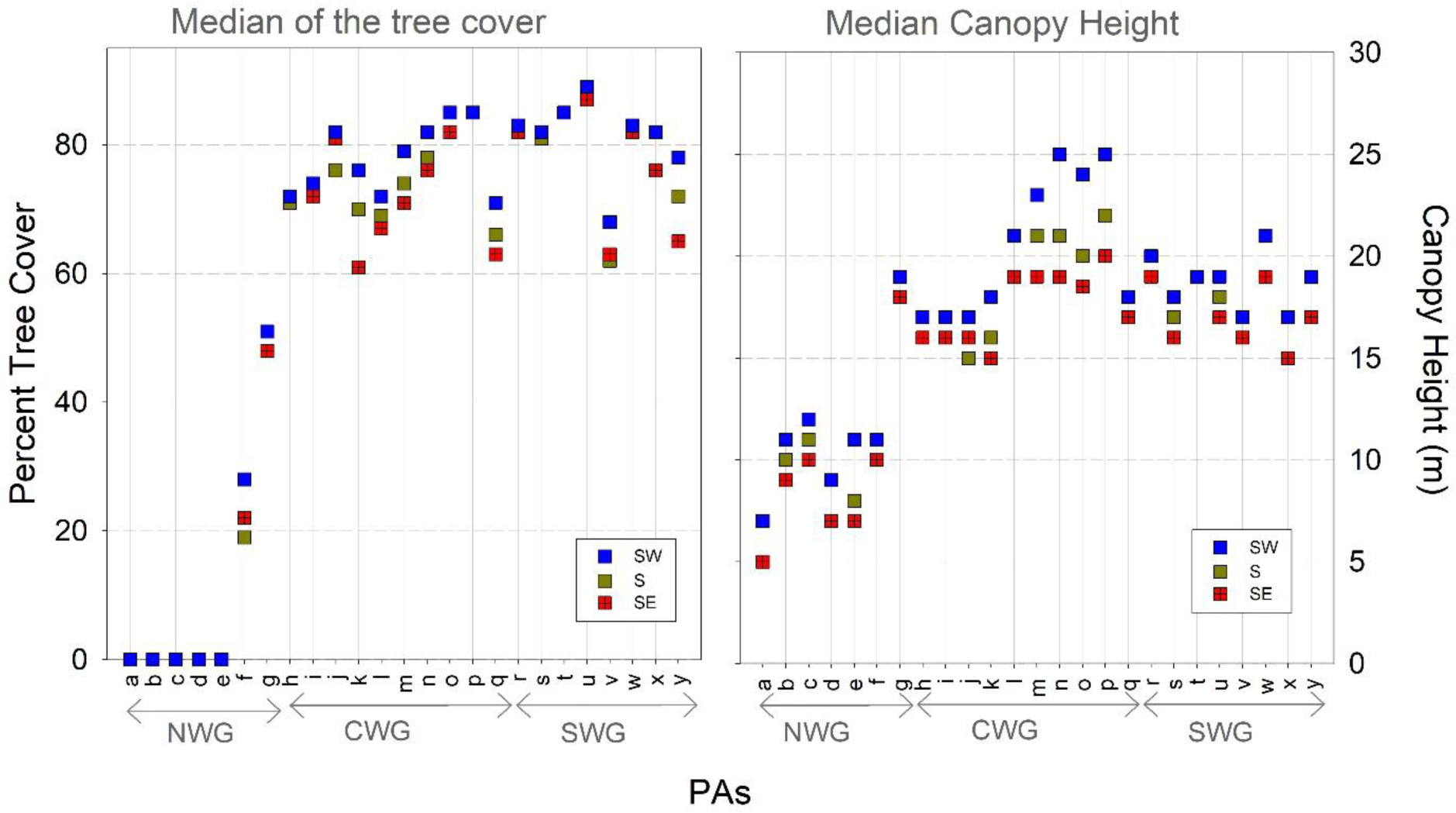
The median TC (left panel) and CH (right panel) on the southwest (SW), south (S) and southeast (SE) aspects in the studied PAs. The abbreviations for the PAs are as in Supplementary Figure S3.

**Supplementary Figure S12.**
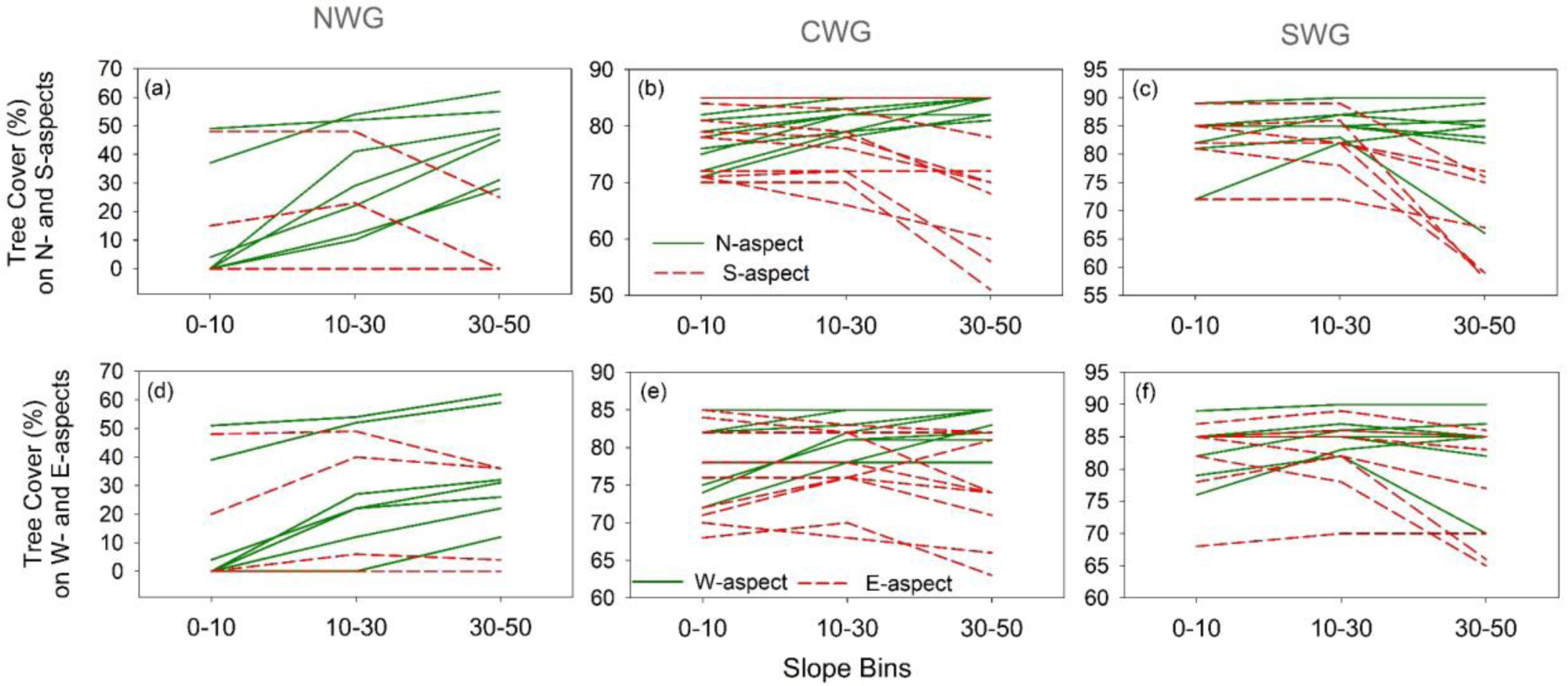
Variation in the TC on N- and S-aspects (a, b, c) and W- and E-aspects (d, e, f) with the slope in NWG (a, d), CWG (b, e) and SWG (c, f). The TC values of S- and E-aspects for the PA Kudremukh (indicated by letter ‘k’ in Suppl. Fig. S3) from CWG and Idukki (indicated by letter ‘v’ in Suppl. Fig. S3) from SWG are removed to reveal the variability in TC in other PAs.

Their values are as below

| PA |  | Aspect | 0°-10° | 10°-30° | 30°-50° |
| --- | --- | --- | --- | --- | --- |
| k | CWG | S | 72 | 71 | 0 |
| k | CWG | E | 70 | 62 | 0 |
| v | SWG | S | 67 | 63 | 0 |
| v | SWG | E | 72 | 64 | 3 |

**Supplementary Figure S13.**
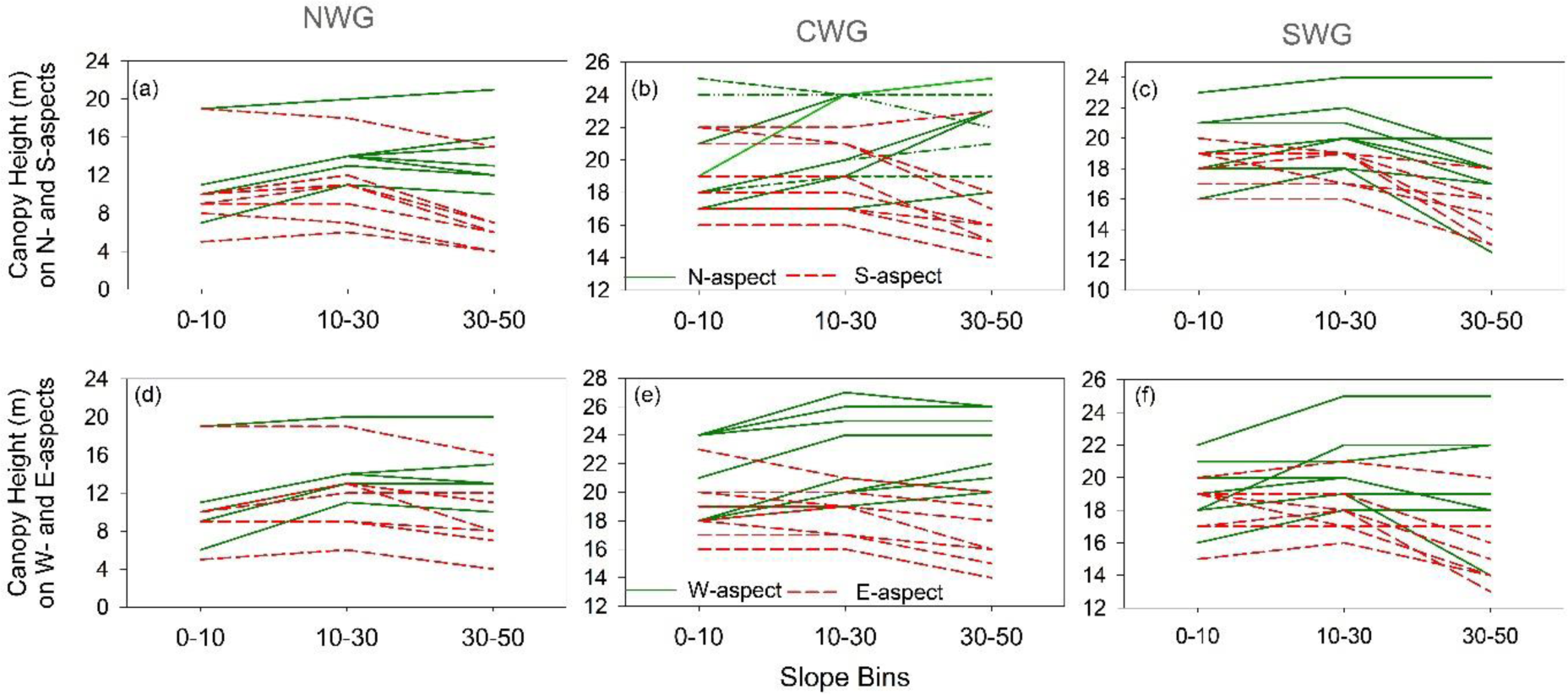
Variation in the CH on N- and S-aspects (a, b, c) and W- and E- aspects (d, e, f) with slope in NWG (a, d), CWG (b, e) and SWG (c, f). The CH values of S- and E-aspects for the PA Kudremukh (indicated by letter ‘k’ in Suppl. Fig. S3) from CWG and Idukki (indicated by letter ‘v’ in Suppl. Fig. S3) from SWG are removed to reveal the variability in TC in other PAs.

Their values are as below

| PA |  | Aspect | 0°-10° | 10°-30° | 30°-50° |
| --- | --- | --- | --- | --- | --- |
| k | CWG | S | 17 | 17 | 5 |
| k | CWG | E | 16 | 14 | 5 |
| v | SWG | S | 16 | 16 | 9 |
| v | SWG | E | 17 | 16 | 7 |

**Supplementary Figure S14.**
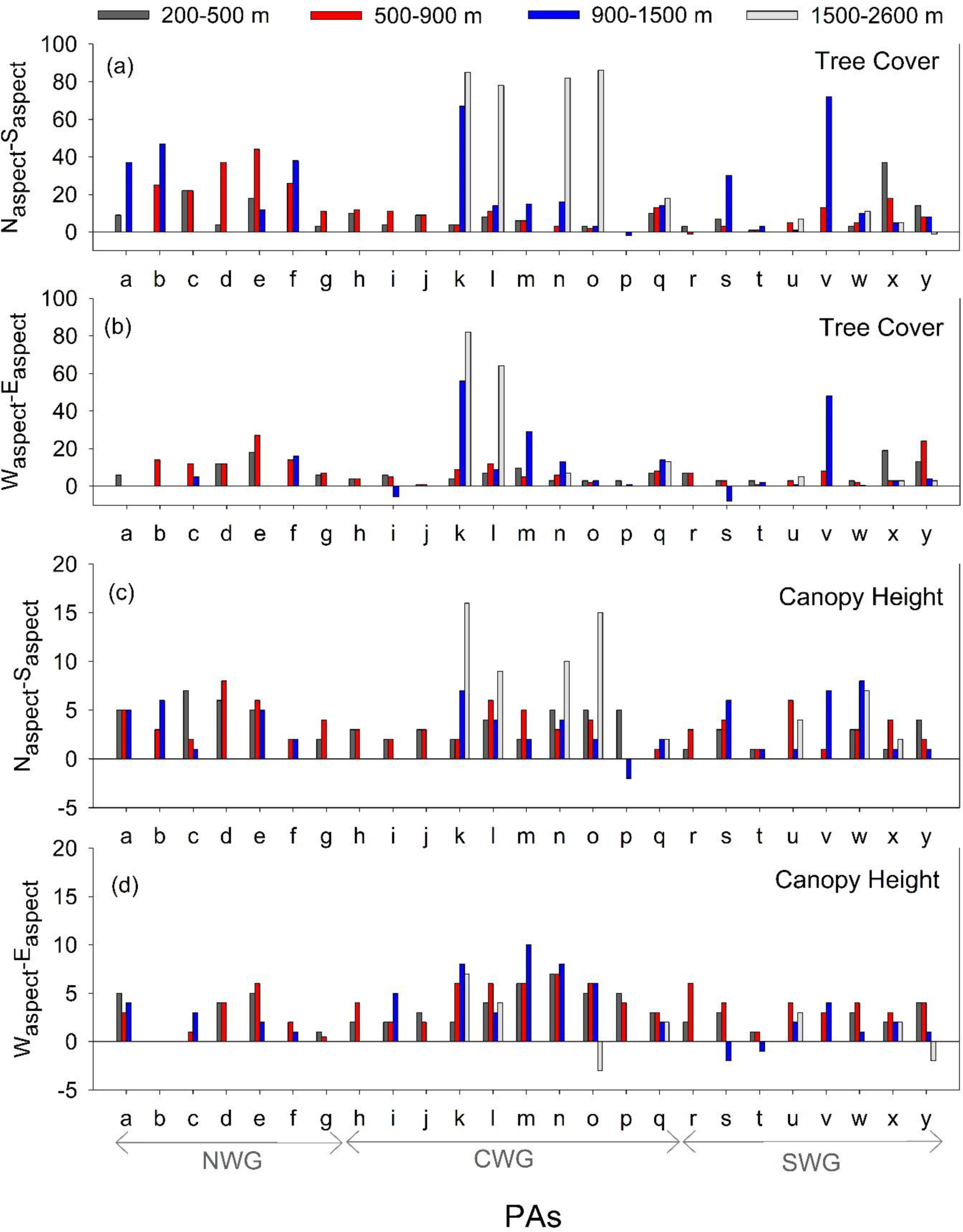
The altitude-dependant variation in the difference between the median tree cover on (a) the N-aspect and S-aspect and (b) the W-aspect and E-aspect. The same for canopy height is shown in (c) and (d). The abbreviations for the PAs are as in Supplementary Figure S3.

**Supplementary Figure S15.**
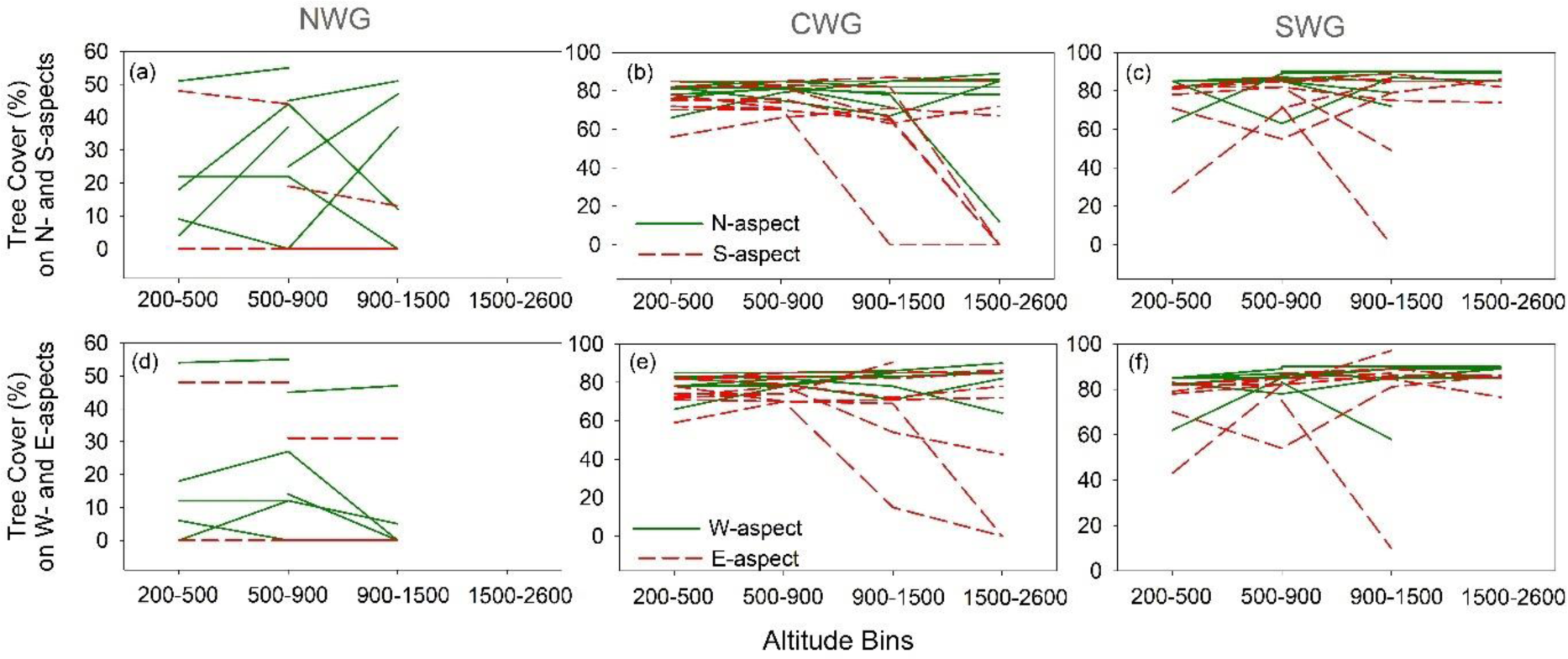
Variation in the TC on N- and S-aspects (a, b, c) and W- and E-aspects (d, e, f) with altitude in NWG (a, d), CWG (b, e) and SWG (c, f).

**Supplementary Figure S16.**
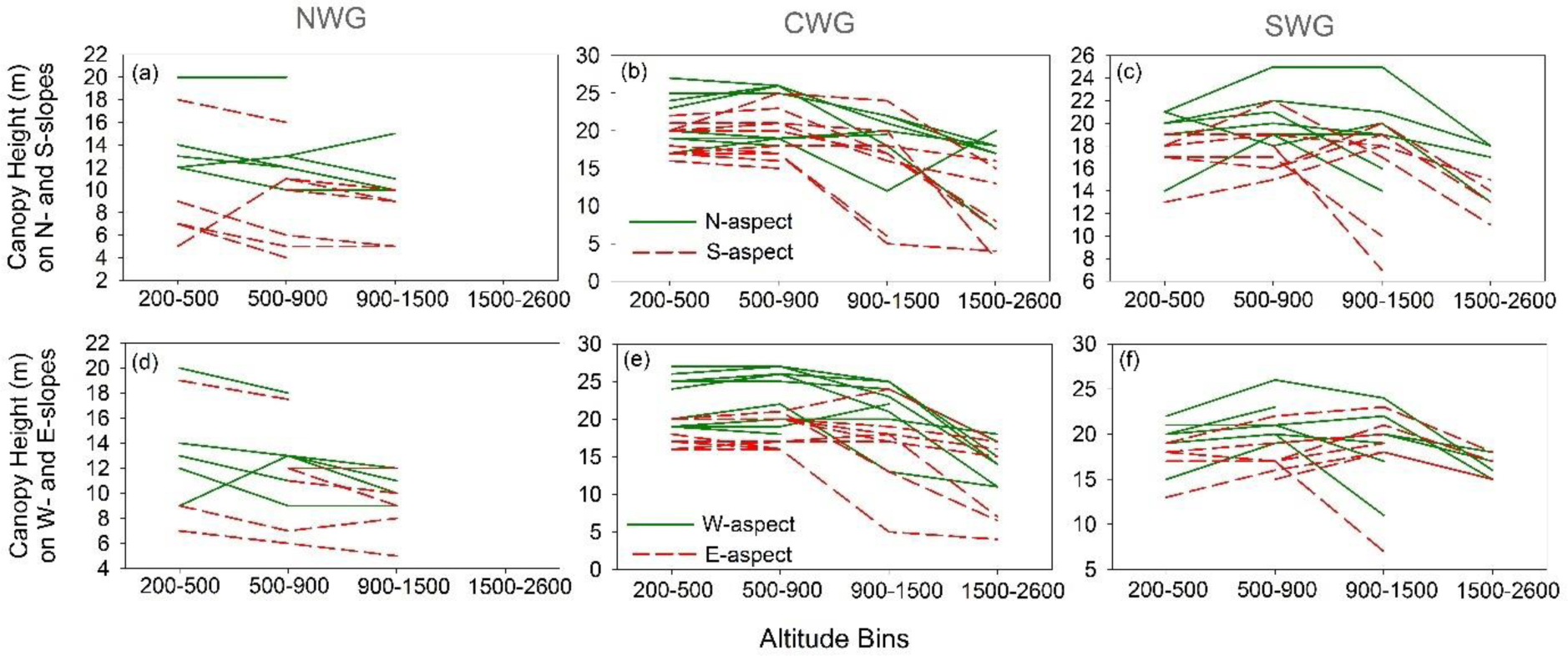
Variation in the CH on N- and S-aspects (a, b, c) and W- and E-aspects (d, e, f) with altitude in NWG (a, d), CWG (b, e) and SWG (c, f).

